# Accurate Protein Domain Structure Annotation with DomainMapper

**DOI:** 10.1101/2022.03.19.484986

**Authors:** Edgar Manriquez-Sandoval, Stephen D. Fried

## Abstract

Automated domain annotation plays a number of important roles in structural informatics and typically involves searching query sequences against Hidden Markov Model (HMM) profiles. This process can be ambiguous or inaccurate when proteins contain domains with non-contiguous residue ranges, and especially when insertional domains are hosted within them. Here we present DomainMapper, an algorithm that accurately assigns a unique domain structure annotation to any query sequence, including those with complex topologies. We validate our domain assignments using the AlphaFold database and confirm that non-contiguity is pervasive (6.5% of all domains in yeast and 2.5% in human). Using this resource, we find that certain folds have strong propensities to be non-contiguous or insertional across the Tree of Life, likely underlying evolutionary preferences for domain topology. DomainMapper is freely available and can be run as a single command line function.

**HIGHLIGHTS:** DomainMapper generates a unique domain structure annotation, including non-contiguous and insertional domains

Automated annotations of non-contiguous domains are validated against the AlphaFold database

DomainMapper can be easily installed and used by non-experts

Certain folds have strong preferences to be non-contiguous or insertional

**GRAPHICAL ABSTRACT:** 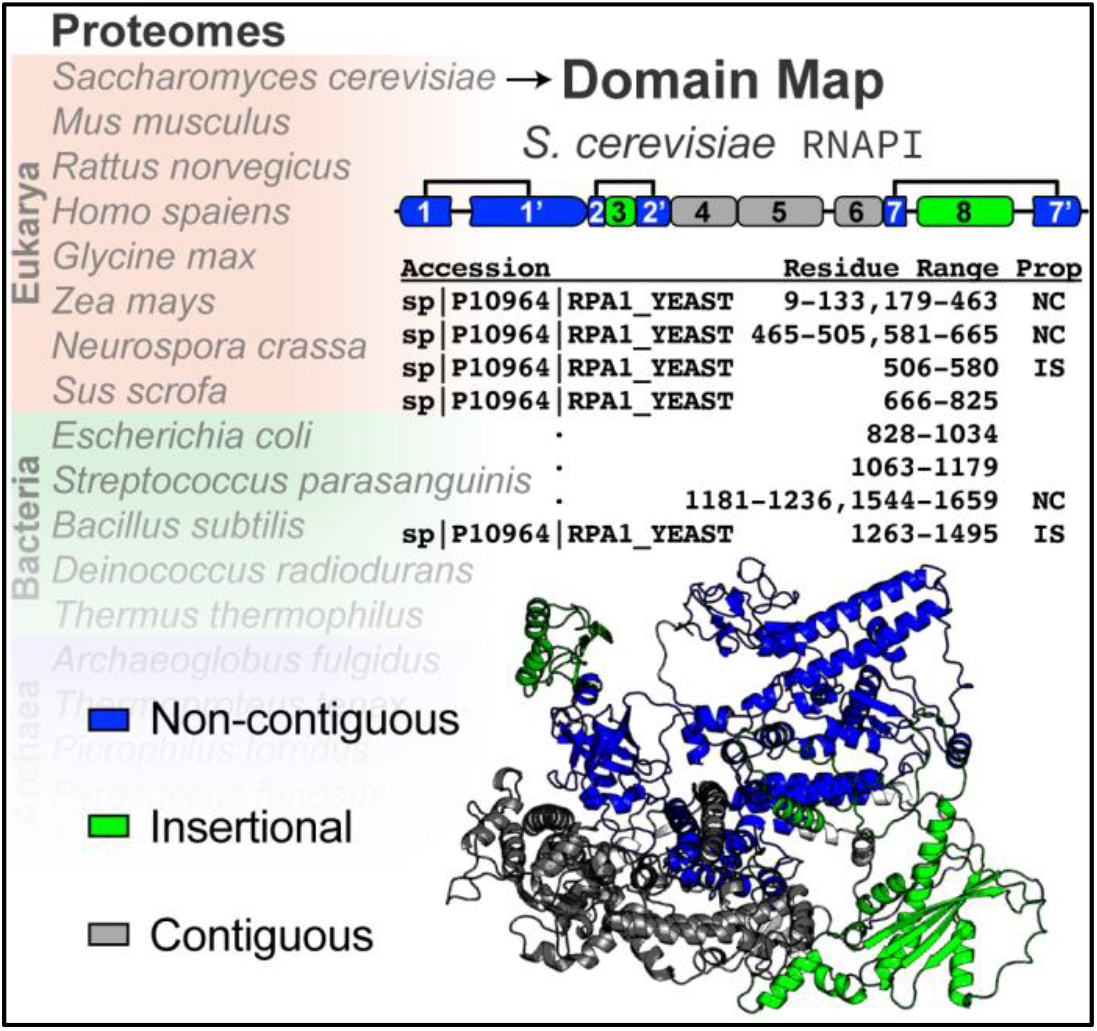

## INTRODUCTION

Proteins that perform complex functions typically comprise of multiple semi-independently folding structural units called domains. These domains can be grouped together into categories that form a ‘periodic table’ of the protein universe, elements that can be merged and combined in myriads of fashions. Automated domain detection has been critical for functional and structural annotation of proteins based on sequence and plays important roles in protein structure prediction (Chothia et al., 2003; Jumper et al., 2021; Vogel et al., 2004), inference of protein function (Friedberg, 2006; Lee et al., 2007; Osadchy and Kolodny, 2011), protein engineering, and rational truncation for protein expression and stabilization.

When using a large database for comprehensive domain annotation (such as SCOP2 (Pandurangan, et al., 2019), CATH (Sillitoe et al., 2019), or Pfam (El-Gebali et al., 2019)), searches will typically yield many matches per protein, of which the majority will be redundant because a given residue range might match to many similar domains with various levels of confidence. Parsers play an important role in reducing this complexity by identifying a ‘minimal’ set of non-overlapping domains such that a given residue range is at most assigned to one domain; where conflicts arise, the ‘best’ match is selected. However, conventional parsers exclude or mis-annotate domains with non-contiguous residue ranges, especially when there are insertional domains lying within them. We initially became intrigued with non-contiguous domains because a recent study (To et al., 2021) found that proteins with non-contiguous domain topologies are typically intrinsically nonrefoldable, suggesting that they lean more heavily on chaperones or other cellular processes to assist their assembly. These findings are consistent with the general understanding that long-range contacts in proteins are more challenging and typically slower to form (Plaxco et al., 1998). Further exploration showed that this feature is not annotated in any of the major databases recording domains, and we reasoned their prevalence may be under-appreciated in structural biology.

ECOD (evolutionary classification of protein domains) (Cheng et al., 2014; Schaeffer et al., 2016) is a relatively new domain classification system that emphasizes distant evolutionary relationships to construct a structural hierarchy and has been particularly useful for computational structure prediction (Jones and Kandathil, 2018) and ancestral protein analysis (Longo et al., 2020). We therefore set out to use the ECOD system as a starting point to create a program called DomainMapper, an algorithm that can accurately predict a unique highest-confidence domain map for any protein based on sequence. Importantly, DomainMapper accurately assigns residue-ranges to domains with non-trivial topologies, specifically non-contiguous domains, insertional domains, and circularly permuted domains. We demonstrate the accuracy and generality of our algorithm by comparing sequence-derived predictions for domain ranges against the AlphaFold database. Using this tool, we have documented the domain structures for the proteomes of seven model organisms, including humans. These annotations enable us to see clear trends of particular folds being significantly more (or less) amenable to non-contiguous topologies over others, patterns that recapitulate underlying biophysical and evolutionary aspects of these structural units. We expect that the accurate domain maps described here, and the ease of use of our algorithm, will serve as a resource for the structural biology community.

## RESULTS AND DISCUSSION

### Accurate Prediction of Non-contiguous Domains with DomainMapper

Following installation (see Implementation and STAR Methods), DomainMapper is supplied with an hmmscan (Eddy, 1996; Eddy, 2011) output file, and generates a unique maximum-confidence domain map for each protein. Each domain is assigned a residue range (that is possibly non-contiguous) and is described by a set of labels in the ECOD hierarchy, specifically, an architecture (e.g., a/b three-layer sandwich), an X-group (similar to a fold, e.g., P-loop NTP hydrolase), a T-group (similar to a superfamily), and a F-group (a more specific classification tied to a particular function). Proteins can be assigned multiple domains, which themselves can be assigned several different topological annotations. Most domains are contiguous and do not get assigned any labels, however some domains can be labeled: (i) non-contiguous (NC); (ii) circular permutant (CP); (iii) insertional; (iv) non-contiguous and circular permutant; (v) insertional and circular permutant; or (vi) non-contiguous and insertional. We find that non-contiguous domains (and insertional domains) are universal (Figure 1A). In humans, 2.49% of all domains have a non-contiguous residue range (N = 3,985). In yeast, this fraction rises to 6.54% (N = 479). Insertional domains are generally 4–5 times less frequent (0.62% of all domains in humans (N = 998), 1.27% in yeast (N = 93)). This is because non-contiguous domains are interrupted by disordered “linker” regions more often than they are interrupted by other structured elements.

**Figure 1.**
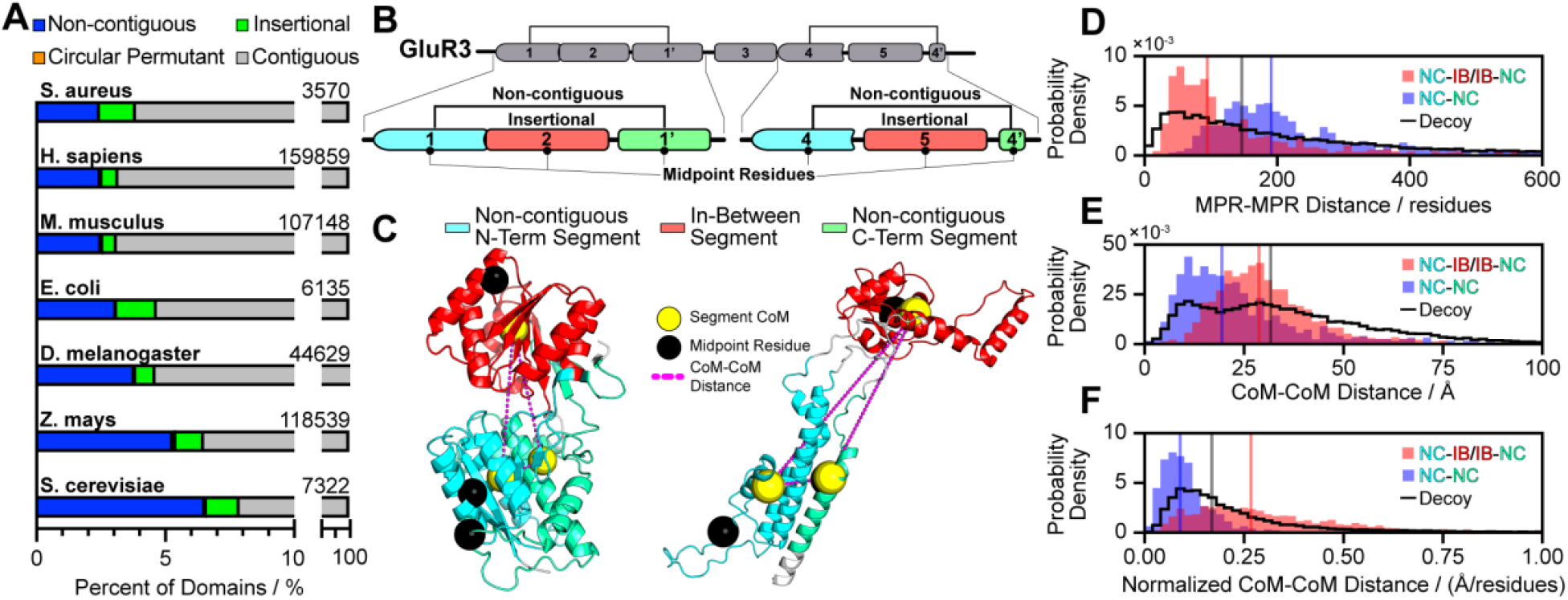
Comprehensive Accurate Annotation of Protein Topology with DomainMapper. (A) Bar charts showing the domain compositions of seven model proteomes, and the frequency of topological labels assigned to individual domains by DomainMapper: non-contiguous (NC, blue); circular permutant (CP, orange); insertional (IS, green); and contiguous (CON, gray). *S. aureus*, N = 3,570 (NC: 2.46%, CP: 0.00%, IS: 1.40%, CON: 96.14%); *H. sapiens*, N = 159,859 (NC: 2.49%, CP: 0.07%, IS: 0.62%, CON: 96.82%); *M. musculus*, N = 107,148 (NC: 2.52%, CP: 0.07%, IS: 0.52%, CON: 96.89%); *E. coli*, N = 6135 (NC: 3.10%, CP: 0.02%, IS: 1.55%, CON: 95.33%); *D. melanogaster*, N = 44,629 (NC: 3.81%, CP: 0.06%, IS: 0.73%, CON: 95.40%); *Z. mays*, N = 118,539 (NC: 5.30%, CP: 0.12%, IS: 1.08%, CON: 93.50%); *S. cerevisiae*, N = 7,322 (NC: 6.54%, CP: 0.07%, IS: 1.27%, CON: 92.12%). (B) Domain map of human glutamate receptor 3 (GluR3, Uniprot: P42263) with five distinct domains: a single contiguous domain (3), two non-contiguous domains (1 and 4) and two insertional domains (2 and 5). Shown is a detailed view of the non-contiguous domains (with N-terminal and C-terminal segments in cyan and teal, respectively) and the insertional domains they enclose in red. Black points mark the midpoint residue (MPR) for each segment. (C) AlphaFold structures of the non-contiguous ranges shown in panel B. Regions in cyan, red, and green correspond to the residue ranges of the three indicated segment types from the DomainMap. Each segment contains two spheres that represent the 3-D coordinates of the MPR (black) and center of mass (CoM, yellow). The magenta dashed lines represent the distance between CoMs. (D-E) Distribution of the distances between segment (D) MPRs and (E) CoMs of all non-contiguous domains in *S. cerevisiae* (N = 479). In blue, distances between N-terminal and C-terminal non-contiguous segments; in red, distances between the in-between (IB) segment and the two flanking non-contiguous segments. The distributions shown as black lines represent distances between decoy “pseudo-NC” segments constructed by splitting all the contiguous domains in *S. cerevisiae* (N = 6,745) and performing the same analysis. (F) As in Panel E, except that all CoM distances are normalized by their corresponding MPR distances for each pair of segments. Distributions use the same color scheme as above.

In the case of a non-contiguous domain, residue segments that come together close in space may be far in sequence (Figure 1B-C), due to an interruption caused by a separate domain, linker, or disordered region. DomainMapper can accurately ‘connect’ these two segments together under the heading of the same domain. For instance, in the structure of human glutamate receptor (Uniprot: P42263, Figure 1B-C, left), we find one periplasmic binding domain (#1, cyan and teal) becomes interrupted with a second periplasmic binding domain (#2, red). We assigned the residue ranges to these two domains based on sequence alone, though to illustrate their accuracy we have painted those ranges onto the AlphaFold structural prediction for GluR3 (Figure 1C, left). From visual inspection, one can see the centers of mass (CoMs, yellow spheres) of the two segments assigned to the non-contiguous domain are much closer in space to each other than either of them are to the center of mass of the ‘in-between (IB) segment,’ even though both NC segments are closer to the in-between segment in sequence space (which can be quantified as the difference between the ‘midpoint residue’ (MPR, black spheres) of each region (Figure 1C). For this analysis, we calculate the IB segment as all residues enveloped by the NC segments, as not all non-contiguous domains are interrupted by structured insertional domains.

Leveraging the AlphaFold database of protein structure predictions, we can assess the generality of our approach by surveying 479 non-contiguous domains within the *S. cerevisiae* proteome (Figure 1D-F) and compare our prediction for the domain boundaries against the predicted structure. For each non-contiguous domain, we located the center of mass (CoM) and midpoint residue (MPR) for the segments that make up the non-contiguous (NC) domain as well as those of the regions that lie in-between (IB). Unsurprisingly, the MPR-to-MPR distance for NC-IB pairs (red) are generally closer to each other than the MPR-to-MPR distance for NC-NC pairs (blue), with median distances of 94.50 residues and 191.0 residues, respectively (Figure 1D). On the other hand, if we compare CoM-to-CoM distances (Figure 1E), it is apparent that NC-NC CoM distances are much shorter than NC-IB CoM distances, with the median distance of 19.41 Å and 28.76 Å, respectively. To further test DomainMapper’s non-contiguous domain residue range annotations, we sought to compare the NC-NC CoM distribution (blue) to a decoy set (black line), in which contiguous domains were split into halves, and CoMs and MPRs were assigned to the two ‘pseudo-NC’ segments of the halves. Satisfyingly, we found that CoM-to-CoM distances were much shorter among actual NC domains (median, 19.41 Å) compared to decoys (median, 31.69 Å). These trends are even more apparent when we normalize Euclidean (3-D) distances with the residue (1-D) distances (i.e., CoM-to-CoM distance divided by MPR-to-MPR distance; Figure 1F), a metric that adjusts for the inherent tendency of segments that are close in sequence to be close in space. We find that the distribution of these normalized distances for NC-NC (blue) pairings are significantly shorter and less variable than NC-IB (red) pairings (Figure 1F, P < 10^−153^ by the Mann-Whitney rank-sum test).

To further demonstrate the accuracy of DomainMapper’s non-contiguous domain residue range annotations, we calculated the radii of gyration (normalized to domain length, *R*_*g*_/*N*) of all non-contiguous domains in *S. cerevisiae* and compared them to the *R*_*g*_/*N* of contiguous domains. Accurate residue ranges for NC domains, we hypothesized, would generate a distribution of *R*_*g*_/*N* comparable to that of contiguous domains. Strikingly, we found that *R*_*g*_/*N* is even smaller on average for NC domains (median 0.088 Å/residues) than it is for contiguous domains (median 0.123 Å/residues; P < 10^−37^ by the Mann-Whitney rank-sum test; Figure S1A). Hence, we conclude that DomainMapper accurately annotates non-contiguous domains and that in general non-contiguous domains are inherently more structurally compact than contiguous domains.

### DomainMapper Captures Cases with Complex Domain Topology

To assess the generality of our algorithm and challenge it with more complex scenarios, we next sought to test whether non-contiguous domains could be accurately called in the human proteome. Metazoan proteomes typically feature larger proteins, with more domains, oftentimes in the form of repeats, and many more disordered regions (Rebeaud et al., 2021). The human proteome contains 159,859 structural domains in 100,100 proteins (including isoforms), though the frequency of non-contiguous and insertional domains are similar to those of other proteomes (Figure 1A). Again, using the AlphaFold database, we mapped 3,641 domain assignments onto the 1,123 proteins which had AlphaFold structures with non-contiguous properties assigned to them and measured the accuracy of this prediction using the same procedure as above. Again, we found that segments assigned to a single non-contiguous domain were generally accurate, as the CoM-CoM distances for NC-NC (blue) pairings were significantly smaller and less variable than for NC-IB (red) pairings (Figure 2, P = 0 by the Mann-Whitney rank-sum test). Likewise, non-contiguous domains are generally compact, with a median *R*_*g*_/*N* of (0.094 Å/residues) significantly smaller than the median *R*_*g*_/*N* of contiguous domains (0.148 Å/residues; P < 10^−67^ by Mann-Whitney rank-sum test; Figure S1B) – analogous to what we observed in *S. cerevisiae*. These results suggest that DomainMapper generates residue range assignments that lead to compact globular structures, even for the proteins with more complex topologies from metazoan proteomes.

**Figure 2.**
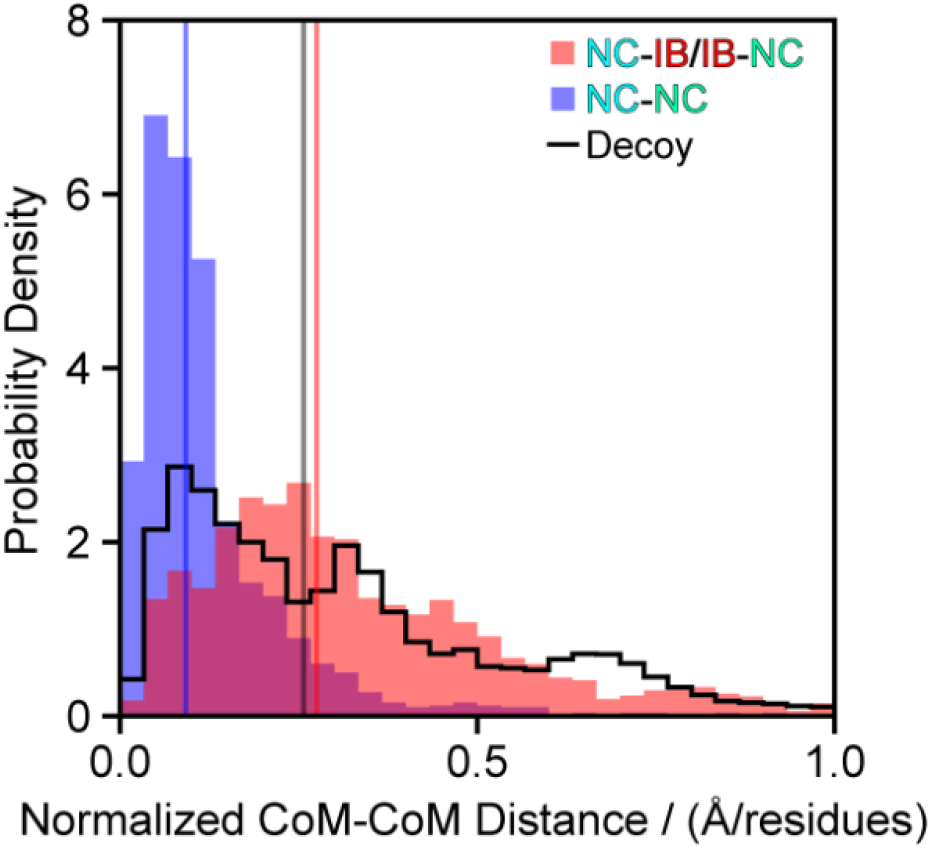
Distribution of the normalized CoM-CoM distances between segments of all non-contiguous domains in *H. sapiens* (N = 3,641) and decoys generated from all contiguous domains from *H. sapiens Chr. 6* (N = 15,854) with AlphaFold structures. In blue, distances between N-terminal and C-terminal non-contiguous segments; in red, distances between the in-between (IB) segment and the two flanking non-contiguous segments with median values of 0.092 and 0.275 respectively. In black, decoy non-contiguous domains were constructed from all continuous domains in *H. sapiens* chromosome 6 with a median value of 0.257.

As an example to highlight this, we show the automated domain assignment generated by DomainMapper for the β’ subunit of RNA polymerase (RpoC, Uniprot: Q8RQE8 from *Thermus thermophilus*, because its x-ray structure is available (PDB: 6KQG (Li et al., 2020), Figure 3). RpoC has an unusually complex domain structure, consisting of 12 distinct structural domains. Three of these domains are non-contiguous (domain 1 in red/salmon, domain 6 in chartreuse/green and domain 11 in purple/violet) and six are insertional (domains 2, 3, 4, 5, 7, and 12). The algorithm properly recapitulates the complex domain topology of this protein, assembling together the pairs of segments that form the N-terminal domain (1), the central helical domain (11), and the double-psi cradle loop barrel (6).

**Figure 3.**
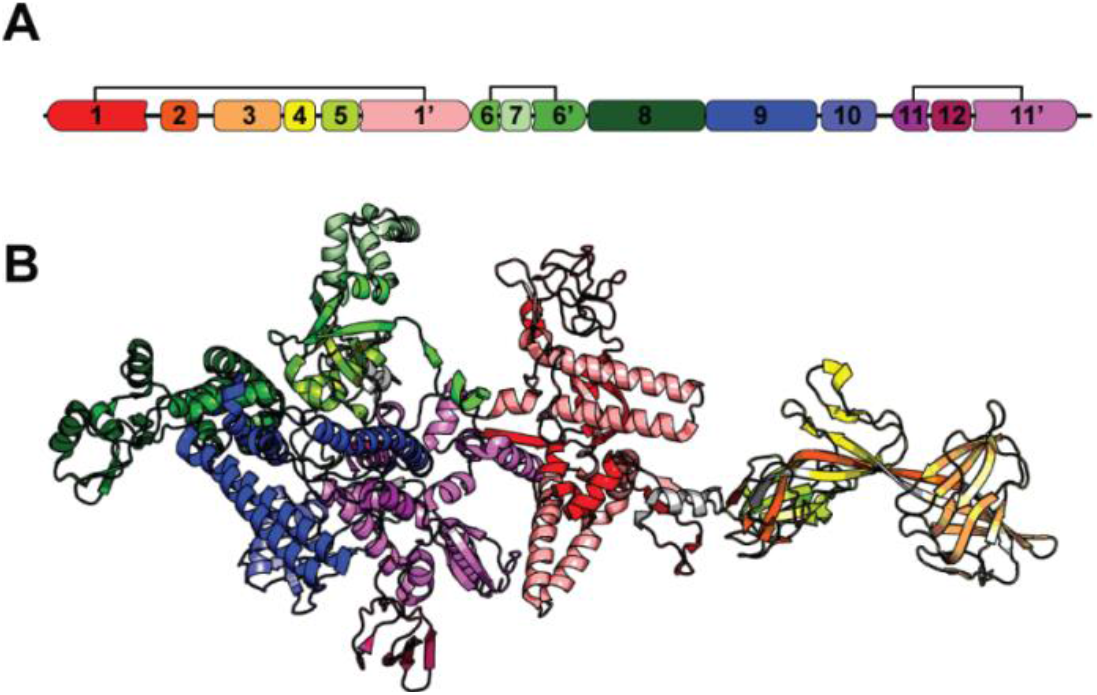
Accurate Annotation of a Complex Domain Topology. (A) DomainMap of *Thermus thermophilus* β’ subunit of RNA polymerase (RpoC, Uniprot: Q8RQE8) based on sequence alone. (B) Projection of the domain assignments from panel A onto the RpoC x-ray crystal structure (PDB: 6KQG). Regions without domain assignments are shown in light gray.

### Non-contiguity Is Systematically Enriched in Different Folds

In the ECOD formalism, all domains are assigned a position in a hierarchy entirely based on evolutionary relationships, termed X-groups, H-groups, T-groups, and F-groups. These classifications correspond loosely to folds (X-groups), superfamilies (T-groups), and families (F-groups) from the SCOP classification system. Using this formalism, we counted the number of times domains in a given X-group had a non-standard topology (e.g., non-contiguous, insertional, or circularly permuted), and discovered that these properties are unevenly spread across protein domain space (Figure 4).

**Figure 4.**
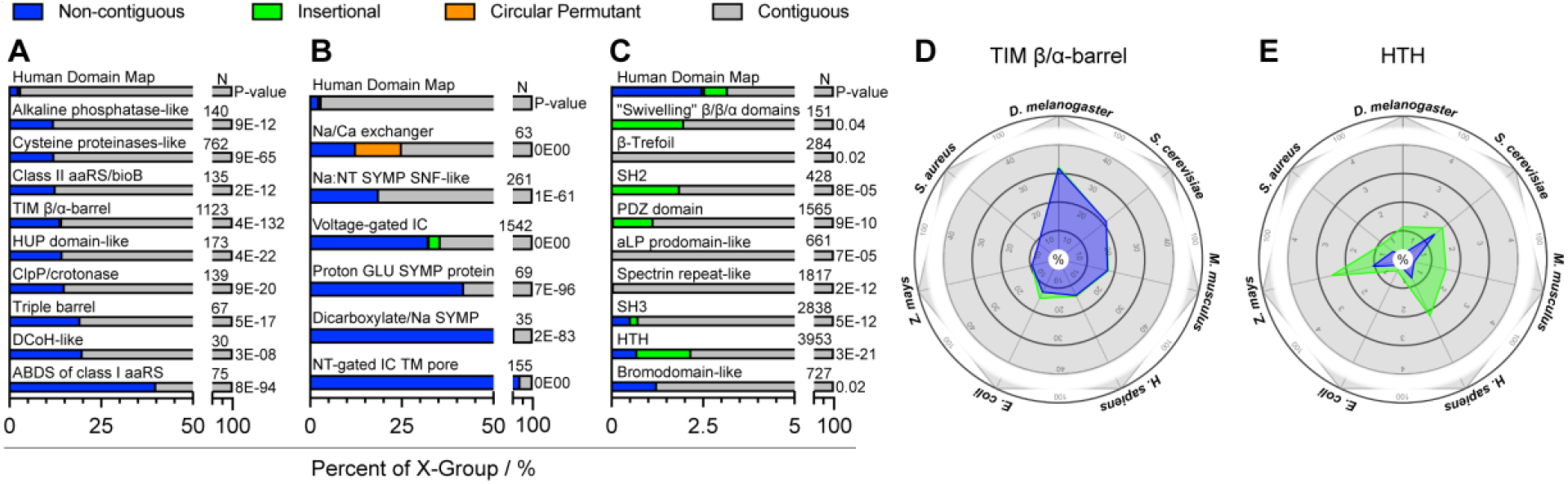
Non-contiguity correlates with particular fold-types. (A) Bar charts showing the frequency of non-contiguous (blue), circular permutant (orange), insertional (green), and contiguous (gray) domains across the human proteome for several X-groups that are enriched for non-contiguity relative to other human protein domains. Also given are the number of domains in that X-group identified in total (N), and the P-value by the chi-square test that the over-/under-representation of these labels within the given X-group, relative to the overall human proteome, are not due to chance. (B) Analogous to panel A, but for several specialized X-groups related to transporters that are ultra-enriched for non-contiguity (C) Analogous to panel A, but for several X-groups that are de-enriched for non-contiguity relative to human protein domains. (D) Polar chart showing the frequency of non-contiguous (blue), and insertional (green) domains across TIM barrels in seven species. (E) Polar chart showing the frequency of non-contiguous (blue), and insertional (green) domains across helix-turn-helix domains in seven species.

Using the human proteome as a model, we found amongst the universal folds, TIM barrels, alkaline phosphatase-like folds, and ClpP/crotonase domains are significantly more likely to host inserts (i.e., be non-contiguous): specifically, 6-fold, 5-fold, and 6-fold more frequently than average (2.49%) (Figure 4A). These enrichments are highly statistically significant in relation to their expected frequencies based on the chi-square test (P-values of 4 × 10^−132^, 9 × 10^−12^, and 9 × 10^−20^, respectively). All of these folds have α/β architectures, and from biophysical studies, TIM barrels and alkaline phosphatase are generally noted for their great stability (Dong and Zeikus, 1998; Sterner and Höcker, 2005; Romero-Romero et al., 2015; Zees et al., 2009). It would seem that this stability could have served as an important foundation to make these folds more amenable to accommodating large insertions within their domain.

Though less universal, we also discovered an unusually strong preference for folds associated with aminoacyl-tRNA synthetases (aaRS) to be non-contiguous (Figure 4A). This corresponds to the X-groups for the class II aaRS core fold, HUP domains (which contain the core class I aaRS fold), and anticodon binding domains. For these categories, non-contiguity is enriched 5-fold, 6-fold, and 16-fold respectively. These observations lend support to the view that vectorial protein synthesis by translation could play an important role in assisting non-contiguous domains from properly folding (and preventing non-native inter-domain contacts from forming) (Frydman et al., 1999; Liu et al., 2019). Hence, the emergence of translation and the synthetases probably occurred together to enable the more generation of more complex domain topologies, including – ironically – the synthetases themselves (Fried et al., 2022).

We found that the more specialized X-groups involved in transporters are hyper-enriched for non-contiguity (Figure 4B) (Coyote-Maestas et al., 2021), including neurotransmitter-gated ion-channel folds (28-fold), Proton glutamate symport folds (16-fold), and voltage-gated ion channels (13-fold). These X-groups are highly attested in the human proteome for which recent gene duplication and many paralogs are available. Hence, these results suggest that non-contiguous topologies could play a pervasive role in enabling signal-recognizing domains to communicate allosterically with transmembrane-regions during active transport.

In addition to looking for enrichments, we also sought to characterize folds that appear to be systematically depleted of non-contiguity (Figure 4C). Amongst universal folds, this cohort is enriched for helix-turn-helices (HTH, also known as three-helix bundles), SH3 domains, α-lytic protease prodomain-like, PDZ domains, and the SH2 domains. The first two groups are 4 and 5-fold *de*-enriched for non-contiguous domains relative to expected frequencies and the latter three are never non-contiguous (with P-values according to the chi-square test of 3 × 10^−21^, 5 × 10^−12^, 7 × 10^−5^, 8 × 10^−10^ and 6 × 10^−5^ respectively). These X-groups have several features in common that make them nearly the opposite of the folds that prefer non-contiguity: they are mostly small (<100 amino acids), and they are primarily all-β or all-α, but never α/β. In some cases, these folds are known to have more modest stability and to fold quickly (e.g., HTH and SH3; Religa et al., 2007; Grantcharova and Baker, 1997; Viguera et al., 1994). These preferences seem explainable, as rapid folding of an insertional domain during co-translational folding could help reduce its interference with the folding of the straddling non-contiguous segments. X-groups that are deprived of non-contiguous topologies are, in many cases, enriched to be insertional (e.g., PDZ domains, SH2 domains, and HTH), suggesting that these topological statuses anti-correlate with one another. Hence, we can conclude that stability and architecture play important roles in separating which proteins can easily support (or not) non-contiguity.

The trends that we have described here, though based on the human proteome, are general across numerous species that we have mapped domains for (Figure 4D-E). For instance, in all organisms that we have surveyed, TIM barrels are more likely to have non-contiguous topologies (Figure 4D), whereas helix-turn-helices are less likely to have non-contiguous topologies (Figure 4E). These observations suggest that these different preferences in topology are inherent in the physical qualities of these different protein folds, irrespective of species.

### Certain Folds Are Enriched as Insertional Domains

We also carried out an analysis to determine how the insertional status is distributed across the domains used in the human proteome. In general, insertional domains are comparatively rarer (∼1%; true in all species, cf. Figure 1A), implying that it is generally more straight-forward to create multi-domain proteins by appending (or prepending) domains to a protein in a tail-to-head manner or interrupt non-contiguous domains with disordered linkers. That said, certain X-groups appear more amenable to be inserted (Figure 5A), notably: cradle loop barrels, FKBP-like domains, and rubredoxin-like domains (for which the insertional status is enriched 7-fold, 16-fold, and 12-fold respectively). These X-groups contain relatively small folds (≤ 70 residues) with N- and C-termini facing in similar directions, which would minimize disruption to the host non-contiguous domains. Both cradle loop barrels and rubredoxin-like domains have a strong preponderance to insert into domains associated with class I aminoacyl-tRNA synthetases (Table S1). It has been suggested that cradle loop barrels share an evolutionary common origin with nucleic acid-binding proteins (Ammelburg et al., 2007), consistent with an early event in which it inserted into an ancestral class I tRNA synthetase domain, and proceeded to co-evolve as a two-domain cassette. Insertion of cradle loop barrels did not occur exclusive into synthetase families (called F-groups in the ECOD system); insertion also occurred into M18 aminopeptidase, pyruvate kinase, and ubiquitin E1 families. Likewise, rubredoxins were also found to insert into SIR2 deacetylase domains and ubiquitin E3 families (see Table S1).

**Figure 5.**
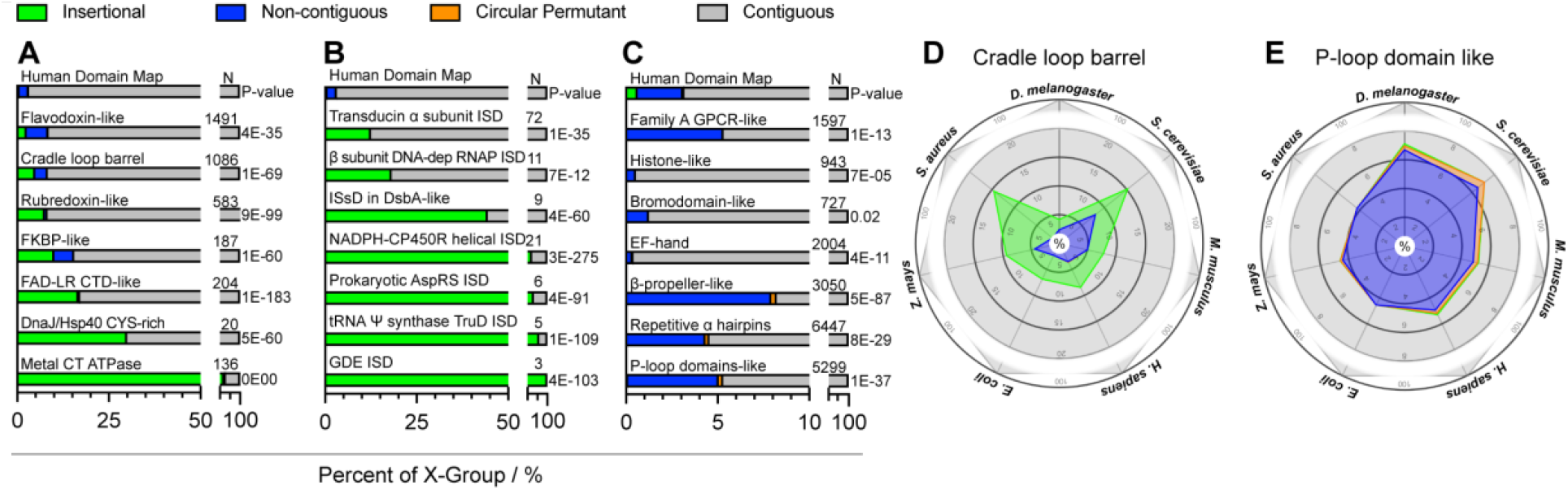
Insertionality correlates with particular fold-types. (A) Bar charts showing the frequency of non-contiguous (blue), circular permutant (orange), insertional (green), and contiguous (gray) domains across the human proteome for several X-groups that are enriched to be insertional relative to other human protein domains. Also given are the number of domains in that X-group identified in total (N), and the P-value by the chi-square test that the over- /under-representation of these labels within the given X-group, relative to the overall human proteome, are not due to chance. (B) Analogous to panel A, but for several X-groups that are de-enriched to be insertional relative to human protein domains. (C) Analogous to panel A, but for several specialized X-groups that are ultra-enriched to be insertional (Note that ISD stands for ‘insertional domain’). (D) Polar chart showing the frequency of non-contiguous (blue), and insertional (green) domains across cradle loop barrels in seven species. (E) Polar chart showing the frequency of non-contiguous (blue), and insertional (green) domains across P-loop NTPase domains in seven species.

Tellingly, DomainMapper found a handful of X-groups that are ultra-enriched to be inserted (with some over 50%), and satisfyingly all of these fold-types are annotated as an insertional domain (ISD, Figure 5B), implying that DomainMapper recapitulates this annotation. In these cases, it is likely that the domain in question arose originally as an insert within a host domain, and that the two-domain cassette served as a common ancestor for further appearances of the inserted domain. We also encountered several folds that are systematically deprived of being insertional (Figure 5C). Some of these are small all-α domains that are quite extended (i.e., where the N- and C-termini point away from each other), which would readily explain why they would be difficult to insert on geometrical grounds (notably, the EF-hand, histone-like, and repetitive α-hairpins). We also found that P-loop NTPase domains and β-propeller domains virtually never occur as inserts (P-values of 1×10^−37^ and 6×10^−87^ by the chi-square test, respectively). Finally, we find that most of the trends described are conserved across species (Figure 5D-E). The cradle loop barrel has a preference to be inserted as does the P-loop NTPase a preference to not be inserted across the seven proteomes we surveyed.

## IMPLEMENTATION

DomainMapper reads the full output file generated from an hmmscan, whose results are organized with the following hierarchy: For each query protein, a table of ‘hits’ describes all the HMM profiles that matched somewhere in the protein sequence. For each hit, a list of ‘HSPs’ (high-scoring pairs) documents all the residue ranges in the query that aligned to the HMM profile. For each HSP, E-values and alignments between HMM and query sequence are given.

Bioinformatically, a non-contiguous domain can be represented in one of two ways: (i) it can be a single HSP wherein the alignment has a large gap in the query sequence; or (ii) it could be broken down into multiple HSPs (Figure 6A). If the latter, the HSPs must necessarily correspond to distinct regions within the HMM.

**Figure 6.**
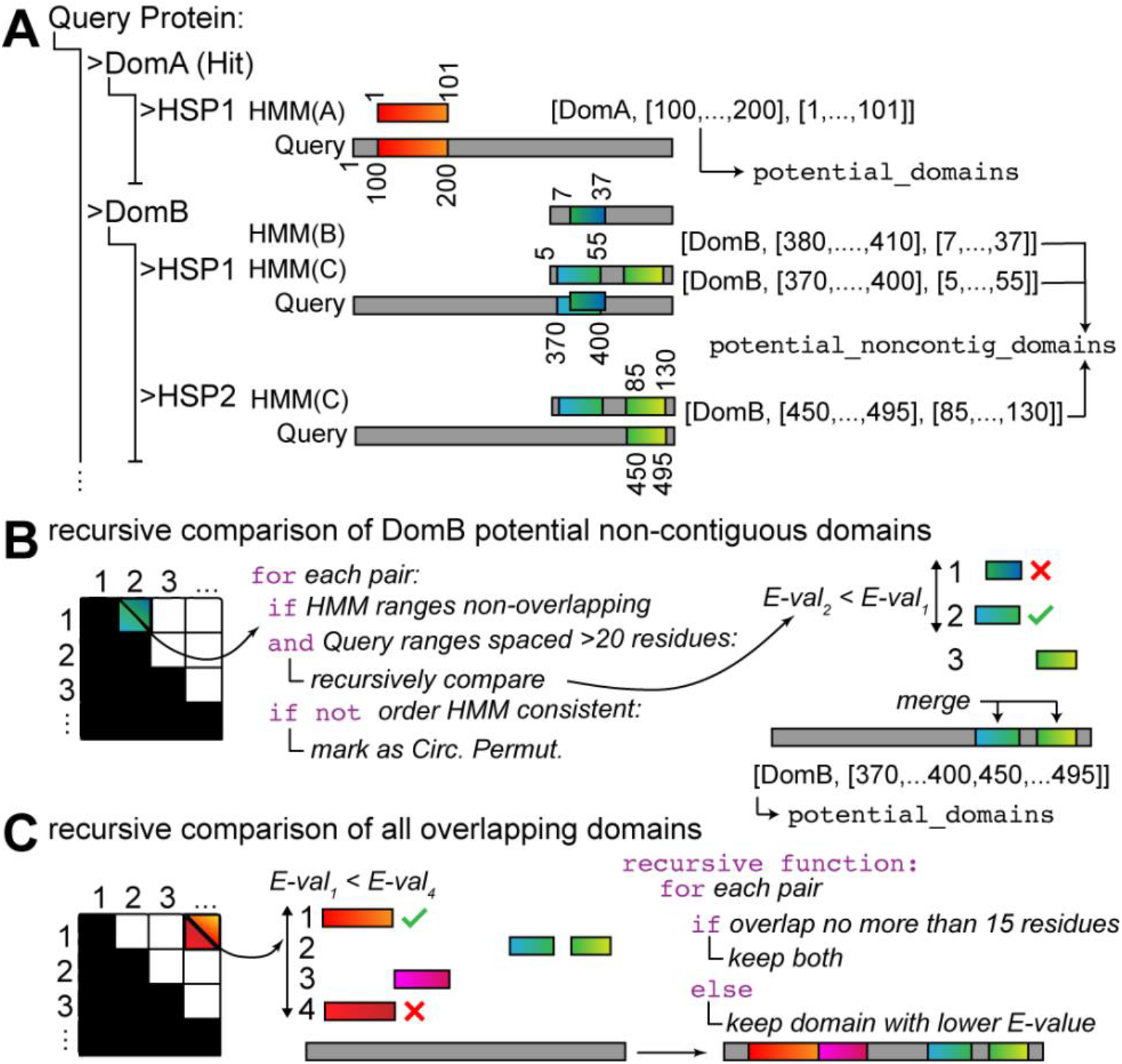
Schematic diagram of DomainMapper’s algorithm. (**A**) Two potential representations of non-contiguous domains. (**B**) Choosing whether to merge or keep separate multiple HSPs (high-scoring pairs) associated with a given hit. (**C**) Choosing the optimal set of potential domains to include in final domain map

To identify structural domains (contiguous or otherwise), DomainMapper uses the following algorithm. For each hit, we ask if there is 1 or >1 HSPs. If there is only 1 HSP, we add that HSP and its residue range to a list of potential domains (assuming its E-value falls below a threshold value of 10^−5^). However, we also check to see if there’s any long gap in the query sequence in the HSP’s alignment (of 35 residues or more, called the intra-HSP gap parameter). If there is, the residue indices of the query sequence corresponding to the gap are removed from the residue range recorded for this potential domain. Importantly, we represent residue ranges as lists of all residue indices, not just as the starting and ending index (e.g., Figure 6A).

If there is >1 HSPs for a hit, we have to consider if such HSPs constitute: (i) separate copies of the domain; (ii) separate segments of a single non-contiguous domain; (iii) a single contiguous domain that simply was divided into separate HSPs by HMMER3 (Figure 6B). All HSPs are recorded along with their residue ranges (deprived of gaps, as before) – though these are not immediately added to the list of potential domains, but instead added to a list of potential non-contiguous domains for this particular hit. Importantly, the residue range of the query and the HMM profile are *both* recorded.

The HSPs are then analyzed in a pairwise manner. If the order of the HSPs is different in the HMM and query, the potential domain is annotated as a circular permutant (CP). HSP pairs which are separated by less than 20 residues (the inter-HSP gap parameter) are then merged together and kept for the next step. If a pair of HSPs’ residue ranges are separated by 20 residues or more in the query, then their residue ranges are retained for potential merging into a non-contiguous domain. Specifically, the retained HSPs that overlap with less than 15 residues (the overlap parameter) are tested recursively into potential mergers ultimately resulting in a set of merged HSPs that maximize query coverage and minimize the combined E-values. These are then added to the list of potential domains. E-values are combined by logarithmic mean.

Finally, we decide which potential domains to keep (Figure 6C). Potential domains are again analyzed in a recursive manner, but now across all hits (domain-types). For each pair, if the two potential domains’ residue ranges overlap (the same overlap parameter applies), then the one with the higher E-value is removed. The potential domains that remain standing after this process of elimination then represent a unique set of high-confidence, non-overlapping, possibly non-contiguous domains for the query protein.

DomainMapper is built to automate this process over all the queries in an hmmscan output file. If domains are being identified in the ECOD structural classification system, the output documents each domain’s architecture (e.g., beta barrels), X-group (e.g., OB-fold), T-group (e.g., Nucleic acid-binding proteins), and family ID (e.g., 2.1.1.118). Otherwise, the domain is documented with just the identifier of the HMM profile.

## USING THE SCRIPT

The python script, DomainMapper.py, is available at https://github.com/FriedLabJHU/DomainMapper and requires BioPython (Cock,P.J.A. et al., 2009) as a dependency. The script should be provided with a HMMER3 hmmscan output file and an output directory path. Sample commands and instructions on running HMMER3 are provided in the README. It will automatically download the latest ECOD domain definitions file and parse it into a binary file. Additionally, sample input and output files for the model organism proteomes are available.

## CONCLUSION

Protein domain annotation is an important part of structural biology, but it has not been standardized. This is both because of the different HMM profile databases available (SCOP2, Pfam, CATH, ECOD, etc.) as well as the semi-arbitrary process of domain parsing. Although Uniprot includes domain annotations for all proteins in its database, these annotations intermingle structural and functional features and do not accurately label domains with complex topologies. Hence, DomainMapper addresses an unmet need in structural biology to provide unique high-confidence structural domain maps in a scalable and flexible manner. Using DomainMapper, we were able to reliably annotate complex topologies such as non-contiguous domains, insertional domains, and circular permutant domains across the Tree of Life. Our analysis shows that certain fold-types are enriched or de-enriched for these topological statuses, observations that correlate with their biophysical properties. Finally, we note that DomainMapper is fast, user-friendly, and user-configurable, which makes it a promising platform to expand the Family & Domains section on Uniprot in the future.

## SUPPLEMENTAL INFORMATION

Supplemental information can be found online at XXX.

## ACKNOWLEDGMENTS

We would like to thank the NSF Division of Molecular and Cellular Biology (2045844) for support. EMS thanks the Program in Molecular Biophysics training grant (NIH T32GM135131). We also thank Liam Longo for advice and suggestions about how to use the ECOD database.

## AUTHOR CONTRIBUTIONS

EMS developed DomainMapper and ran analyses. EMS and SDF discussed the results and wrote the manuscript.

## STAR⋆METHODS

### KEY RESOURCE TABLE

| REAGENT or RESOURCE | SOURCE | IDENTIFIER |
| --- | --- | --- |
| Deposited data |  |  |
| Domain mappings for all proteomes | This paper | 10.5281/zenodo.5931196 |
| All proteomes with canonical and isoforms proteins | This paper | 10.5281/zenodo.5931112 |
| Software and algorithms |  |  |
| HMMER3 v3.3.2 | Eddy S. R., 2011 | http://hmmer.org |
| Python v3.8.6 | Python Software Foundation | https://www.python.org |
| Pymol molecular graphics system | Schrodinger, LLC | https://pymol.org |
| Biopython v1.60 | Biopython.org | https://biopython.org |
| Numpy v1.17.0 | Numpy.org | https://numpy.org |
| SciPy v1.6.1 | SciPy.org | https://scipy.org/ |
| ECOD domain definitions | Cheng et. al., 2014 | http://prodata.swmed.edu/ecod/ |
| DomainMapper 2.0.2 | This paper | 10.5281/zenodo.5930997<br>https://github.com/FriedLabJHU/DomainMapper |
| Other |  |  |
| *Homo sapiens* proteome sequences (canonical and isoforms) Aug. 9<sup>th</sup>, 2021 | Uniprot | https://www.uniprot.org/proteomes/UP000005640 |
| *Escherichia coli* proteome sequences (canonical and isoforms) Aug. 9<sup>th</sup>, 2021 | Uniprot | https://www.uniprot.org/proteomes/UP000000625 |
| *Mus musculus* proteome sequences (canonical and isoforms) Aug. 9<sup>th</sup>, 2021 | Uniprot | https://www.uniprot.org/proteomes/UP000000589 |
| *Saccharomyces cerevisiae* proteome sequences (canonical and isoforms) Aug. 9<sup>th</sup>, 2021 | Uniprot | https://www.uniprot.org/proteomes/UP000002311 |
| *Drosophila melanogaster* proteome sequences (canonical and isoforms) Aug. 9<sup>th</sup>, 2021 | Uniprot | https://www.uniprot.org/proteomes/UP000000803 |
| *Staphylococcus aureus* proteome sequences (canonical and isoforms) Nov. 1<sup>st</sup>, 2021 | Uniprot | https://www.uniprot.org/proteomes/UP000008816 |
| *Zea mays* proteome sequences (canonical and isoforms) Accessed Nov. 1<sup>st</sup>, 2021 | Uniprot | https://www.uniprot.org/proteomes/UP000007305 |
| All Proteome AlphaFold 3D structures | Jumper et al., 2021 | https://alphafold.ebi.ac.uk/download |

## RESOURCE AVAILABILITY

### Lead contact

Further information on the methods, datasets, and computational resources should be directed and will be fulfilled by the lead contact, Stephen D. Fried.

### Materials availability

No new materials were generated in this study.

### Data and code availability

All statistical data reported in this study will be shared by the lead contact upon request. The data reported in this study was generated with DomainMapper v2.0.2, which has been deposited and as of the date of publication is publicly available here: source code (10.5281/zenodo.5930997) and data (10.5281/zenodo.5931196). Future releases of the DomainMapper software will be available at https://github.com/FriedLabJHU/DomainMapper. Domain maps created with DomainMapper v2.0.2, Hidden Markov Model domain alignments created with HMMER3 v3.3.2, and proteome sequence FASTA files accessed from Uniprot on August 9^th^, 2021, and November 1^st^, 2021, have been deposited at (10.5281/zenodo.5931112) and are publicly available as of the date of publication. DOIs are listed in the key resources table. Any additional information required to reanalyze the data reported in this paper is available from the lead contact upon request.

## EXPERIMENTAL MODEL AND SUBJECT DETAILS

Not applicable.

## METHOD DETAILS

### Domain alignments utilizing ECOD domain definitions in HMMER3

The ECOD formalism is not a standard hidden Markov model (HMM) profile available on the HMMER3 webserver and utilization thus requires preparation of the ECOD family HMM profile on a local machine. The prebuilt HMM profiles are available for download from http://prodata.swmed.edu/ecod/ as ecodf.hmm.tar.gz which are regularly updated on a 2–3-month basis. Processing of the ECOD HMM profile was performed in HMMER3 v3.3.2 following a process available in the software documentation summarized here (Eddy, 1996; Eddy, 2011). The extracted ECOD HMM profile download is passed to the `hmmpress` HMMER3 function, which recompresses the profiles into a proprietary binary file and index file that can be utilized to create domain alignments. Domain alignments were performed using the `hmmscan` HMMER3 function with the ECOD binary files and proteome FASTA files as inputs. Since DomainMapper requires the full output from `hmmscan` (as opposed to truncated outputs generated by the `-tblout` and `-domtblout` flags), the `-o` flag was set to true and provided the desired output file name. In the output generated by `hmmscan`, every protein is specified by a query name and Uniprot accession label followed by a summary of the domains detected in the query protein sequence along with their residue ranges. For each hit, a series of residue ranges in the query protein that match it (high-scoring pair, HSP) are provided. For each HSP, an alignment of the query sequence to the HMM model is provided along with several goodness-of-fit metrics (such as score and conditional E-value) and residue ranges, both with respect to the query protein and the HMM. Proteomes were accessed from Uniprot on 9^th^ August 2021 and 1^st^ November 2021 for the seven model organisms reported in this study and aligned in the same way. HMM scanning the 79,038 proteins (isoforms included) of the *Homo sapiens* proteome (Uniprot: UP000005640) took roughly 24 hours on 36 threads on an Intel Xeon W-2295 processor with 18 cores and 64 GB of system memory.

### Mapping of domains

Prior to parsing the HMMER3 output, the latest ECOD domains definitions, which are available at http://prodata.swmed.edu/ecod/asecod.latest.domains.txt, were parsed into a format which allows bottom-up search of the domain hierarchy (F-group, T-group, H-group, X-group, Architecture). This operation is performed upon first install of DomainMapper and reoccurs every 2 months after install to ensure the most up-to-date domain definitions are used. Mapping of domains from the HMMER3 output relies primarily on two metrics, HMM domain coverage and the conditional E-value. In the case of a single domain match to a query range, it is first checked for non-contiguity by searching for gaps between the HMM range and the query range larger than 35 (intra-HSP gap parameter) residues then added to a list of potential domain mappings for that query. In the case of multiple domain assignments, first they are filtered with a permissive E-value cutoff of 1E-2, then they are de-gapped in the same way single domain assignments are (using the intra-HSP gap parameter), combined back together and finally filtered for overlap between adjacent domain assignments fewer than 15 residues (the overlap parameter). The 15-residue overlap is tolerated for cases where domain feathering has occurred, a process wherein throughout the course of evolution one domain “blended” into another with high sequence similarity on terminal ends. Overlapping domains of more than 15 residues (overlap of 70% or more for domains smaller than 30 residues) are compared recursively to find the domain with lowest E-value. These domains are then saved to a list of potential domain mappings for that query. All the domains in the potential mapping list are then checked for circular permutations and cross compared for insertions within other domains. To ensure only high confidence domains are reported in DomainMapper, the last filtering step keeps domains with an E-value cutoff of 1E-5. Lastly, the T-group, H-group, X-group, and Architecture for every mapped domain is found by back searching the F-group in the previously mentioned ECOD domain definitions. This processed is repeated for all query sequences aligned with HMMER3. Mapping of the 157,010 domains from the *Homo sapiens* proteome took roughly 4 minutes on a single thread on an Intel Xeon W-2295 processor with 18 cores and 64 GB of system memory.

### Testing non-contiguous domain mappings

The normalized center-of-mass distance was used to quantify the accuracy of non-contiguous (NC) domain classification by DomainMapper. For all non-contiguous (NC) domains in the Yeast and Human proteomes, the DomainMapper output was parsed for NC domain residue ranges then the centers-of-mass and midpoint residues were located in both the N-terminal and C-terminal non-contiguous segments as well as the interior segment between the non-contiguous segments, the so-called in-between (IB) segment. The normalized center-of-mass distance was defined by the Euclidian distance between centers-of-mass of two segments divided by the absolute difference between the segments’ midpoint residues. A decoy population was also analyzed by taking all contiguous domains and randomly splitting them into pseudo-non-contiguous segments in 30-70% fractions and performing the same functions as with the non-contiguous domains without the in-between analysis. Centers-of-mass were calculated with Biophython (`Bio.PDBParser`) for all segments from AlphaFold 3D structures where available. The Euclidian distance between centers-of-mass and the absolute difference between midpoint residues were calculated with `NumPy` and separated into populations of NC-NC pairs and NC-IB pairs. Radii of gyration were calculated with `NumPy` using the atomic coordinates from structures and atomic masses, except the calculations were conducted on whole domains (either contiguous or non-contiguous) rather than on separate segments. The available AlphaFold structures were downloaded from the EBI bulk download distribution.

## QUANTIFICATION AND STATISTICAL ANALYSIS

All statistical analyses were performed with Python v3.8.6. The chi-square statistics in Figures 4 and 5 were performed with the Python library SciPy v1.6.1 and significance was determined for p-values < 0.05. The total domain counts in Figure 1 are provided by DomainMapper (N = Total Domains) and the total X-group counts in Figures 4A-C and 5A-C were calculated with the Python library NumPy v1.17.0 (N = Total X-groups). All counts and p-values can be found next to their corresponding bars in Figures 1, 4, and 5. The two-sided Mann-Whitney rank sum test in Figures 1D-F and 2 were all performed with SciPy, and significance was determined for p-values < 0.05. Distribution medians in Figures 1D-F and 2 were found using NumPy and are shown as vertical lines for every distribution. P-values for the distributions can be found in the main text and figure captions. No methods were used to determine whether the data met assumptions of the statistical approaches employed.

**Figure S1.**
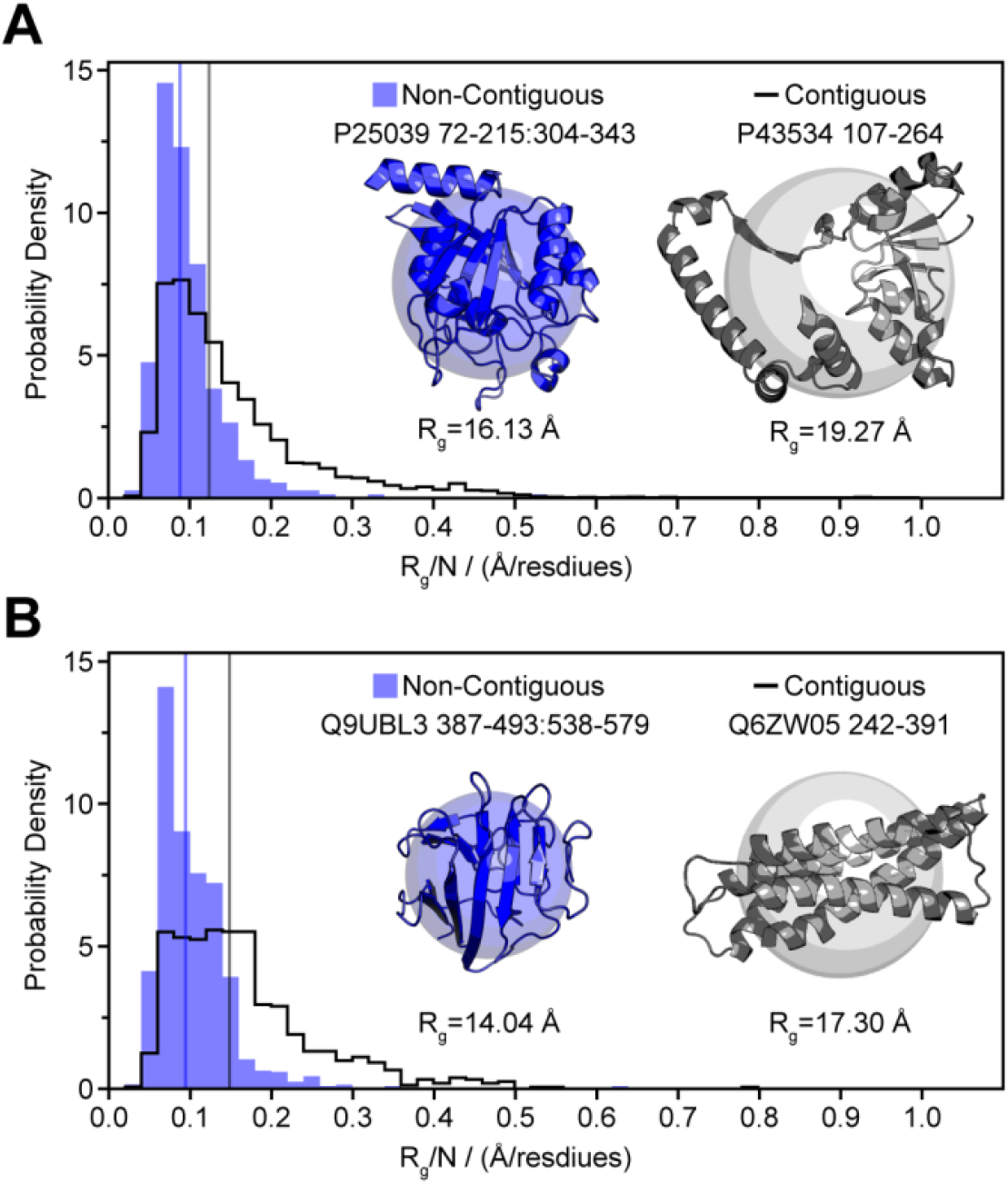
Non-contiguous domains are inherently more compact than contiguous domains, Related to Figure 1 and Figure 2. (A) Distributions for the radii of gyration for all *S. cerevisiae* non-contiguous domains (blue) and contiguous domains (black), normalized by the length of the domain (*R*_*g*_/*N*). The vertical lines represent the medians of the distributions, 0.088 and 0.123 Å/residues, respectively. The two domains pictured on the right are segments of AlphaFold structures chosen to have similar lengths and with *R*_*g*_/*N* representative of their distributions’ median. (B) Similar to panel A but for all *H. sapiens* non-continuous domains (blue) and all contiguous domains from *Chr*. 6 (black).

**Table S1.** Insert-Host F-group pairs of enriched insertional domain X-groups, Related to Figure 5. X-groups and F-groups are shown as they are written in the ECOD database.

| Insertional X-Group | Host X-Group | Insertional F-Group | Host F-Group |
| --- | --- | --- | --- |
| Rubredoxin-like | Rubredoxin-like |  |  |
|  |  | tRNA-synt_1g_1<br>ileS_1st | tRNA-synt_1g |
|  | HUP domain-like |  |  |
|  |  | leuS_1st<br>leuS_2nd | tRNA-synt_1_1st |
|  | SH3 |  |  |
|  |  | KOG0479<br>KOG0482<br>MCM_OB_1<br>MCM_OB_2<br>MCM_OB_4 | TTD_N |
|  | OB-fold |  |  |
|  |  | YciM | MCM_OB |
|  | ETN0001 domain-like |  |  |
|  |  | GCM_2nd | GCM_1st |
|  | Rossmann-like |  |  |
|  |  | KOG1905 | SIR2_1st |
|  | Immunoglobulin-like beta-sandwich |  |  |
|  |  | zf-Sec23_Sec24 | Sec32_BS |
| cradle loop barrel | Rubredoxin-like |  |  |
|  |  | tRNA-synt_1_2nd<br>tRNA-synt_1_2 | tRNA-synt_1g |
|  | HUP domain-like |  |  |
|  |  | tRNA-synt_1_2 | tRNA-synt_1_1st |
|  | Phosphorylase/hydrolase-like |  |  |
|  |  | Peptidase_M18_2nd | Peptidase_M18_1st |
|  | TIM beta/alpha-barrel |  |  |
|  |  | PK_2nd | PK_1st |
|  | Rossmann-like |  |  |
|  |  | E1_FCCH | ThiF_1<br>ThiF_3 |
|  | Catalytic cysteine domain in ubiquitin-activating enzyme |  |  |
|  |  | E1_FCCH | UBA_e1_thiolCys |
| FKBP-like | TIM beta/alpha-barrel |  |  |
|  |  | GH18_chitolectin_chitotriosidase | Glyco_hydro_18 |
|  | TIM beta/alpha-barrel |  |  |
|  |  | GH18_CFLE_spore_hydrolase<br>GH18_SI-CLP | Glyco_hydro_18_1 |

